# Retinoic acid treatment recruits macrophages and increases axonal regeneration after optic nerve injury in the frog *Rana pipiens*

**DOI:** 10.1101/2021.08.04.455100

**Authors:** Valeria De La Rosa-Reyes, Mildred V. Duprey-Díaz, Jonathan M. Blagburn, Rosa E. Blanco

## Abstract

Retinoic acid (RA) plays major roles during nervous system development, and during regeneration of the adult nervous system. We have previously shown that components of the RA signaling pathway are upregulated after optic nerve injury, and that exogenous application of all-trans retinoic acid (ATRA) greatly increases the survival of axotomized retinal ganglion cells (RGCs). The objective of the present study is to investigate the effects of ATRA application on the macrophages in the optic nerve after injury, and to determine whether this affects axonal regeneration. The optic nerve was crushed and treated with PBS, ATRA and/or clodronate-loaded liposomes. Nerves were examined at one and two weeks after axotomy with light microscopy, immunocytochemistry and electron microscopy. ATRA application to the optic nerve caused transient increases in the number of macrophages and microglia one week after injury. The macrophages are consistently labeled with M2-type markers, and have considerable phagocytic activity. ATRA increased ultrastructural features of ongoing phagocytic activity in macrophages at one and two weeks. ATRA treatment also significantly increased the numbers of regenerating GAP-43-labeled axons. Clodronate liposome treatment depleted macrophage numbers by 80%, completely eliminated the ATRA-mediated increase in axonal regeneration, and clodronate treatment alone decreased axonal numbers by 30%. These results suggest that the success of axon regeneration is partially dependent on the presence of debris-phagocytosing macrophages, and that the increases in regeneration caused by ATRA are in part due to their increased numbers. Further studies will examine whether macrophage depletion affects RGC survival.

## Introduction

Retinoic acid (RA) is a vitamin A-derived lipophilic molecule that plays a major role during early development of the nervous system, setting up dorsoventral and anteroposterior patterning of the neural plate and tube [1]. Its later function is to direct the differentiation of various types of neurons and glia by activating the transcription of many genes, including those that encode transcription factors, cell signaling molecules, enzymes and cell surface receptors [1–5].

More recently, it has been found that RA signaling also has a role in the CNS of adult animals, particularly in areas of high neuronal plasticity such as the hippocampus, cortex, and striatum, where cells continue to be generated [1,6–10]. It has also been shown to be involved in the control of rhythms within the brain [11]. In addition, defects in RA signaling may occur in various neurodegenerative diseases including ALS [12, 13] and Alzheimer’s disease [14, 15].

However, of particular relevance to the present study is the growing evidence that RA signaling is reactivated in the adult peripheral nervous system following nerve injury, and thereby enhances the capacity to regenerate. In peripheral nerves, injury causes an upregulation of RA signaling components [16, 17], with one of its receptors, RARβ, being crucial for stimulating neurite outgrowth [18]. However, in mammalian CNS neurons there is no such upregulation of RA signaling after injury, and it was suggested that this, in addition to their inhibitory microenvironment, might be another reason for their poor regenerative capabilities. In contrast, our lab has shown recently that in the frog CNS the RA signaling pathway is indeed upregulated after optic nerve injury, moreover that exogenous application of an RA analogue greatly increases the survival of axotomized retinal ganglion cells (RGCs) [19, 20]. Since RA promotes long-term RGC survival, which generally depends on axons reaching their targets, it is likely that it also promotes the axonal growth of these cells. However, the question of whether RA application stimulates axonal regrowth in the frog remains open, so addressing this question is one of the objectives of this study.

We know that both the numbers and elongation rate of regenerating frog RGC axons can indeed be increased by a single topical application of a growth factor (CNTF or FGF-2) to the optic nerve [21]. The mechanisms for this were not addressed, however, it was subsequently shown that application of these factors also increases the numbers of phagocytic macrophages in the regeneration zone of the injured nerve [22]. Previous work has shown that macrophage infiltration into the eye after lens injury promotes retinal ganglion cell (RGC) survival and regeneration [23–26], and it is now known that RA stimulates macrophage phagocytosis of myelin *in vitro* [27]. These observations lead to the additional questions proposed in the present study: does RA also influence the phagocytic cell populations in the injured optic nerve, and if so, does this affect the regeneration of the axons?

We show that a single application of an RA analogue transiently increases the number of phagocytic macrophages and microglia in the regenerating region of the crushed optic nerve. This treatment also increases the number of regenerating RGC axons, and this increase is prevented by chemical ablation of the macrophages.

## Materials and Methods

### Animals

Adult frogs (*Rana pipiens*) of both sexes were obtained from Connecticut Valley Biological Supply Company (Southampton, MA) and Nasco Worldwide Service to Education, Health, Agriculture, and Industry (Fort Atkinson, WI). They were kept in tanks with recirculating tap water at 19°C. Approximately 50 animals were used for immunohistochemistry and electron microscopy, and 25 more for the axonal regeneration experiments. The study is in accordance with the Guide for the Care and Use of Laboratory Animals of the National Institutes of Health and the recommendations of the Panel on Euthanasia of the American Veterinary Medical Association. Experimental procedures were approved by the Institutional Animal Care and Use Committee of the University of Puerto Rico Medical Sciences Campus.

### Surgical technique for optic nerve crush

The frog was anesthetized by immersion in 0.3% tricaine solution for 10 minutes. To access the optic nerve an incision in the palate was made and then the extraocular muscles of the right eye were retracted. The extracranial portion of the right optic nerve was crushed at the halfway point using forceps (Dumont No. 5). This leaves the meningeal sheath intact but creates a transparent gap that is completely free of axons. We have confirmed the lack of even the smallest of axons in this region by electron microscopic observation; also, crushing in this manner perturbs RGC survival almost as effectively as cutting [28]. A strip of Parafilm was placed under the crushed nerve area to contain any applied solutions. The palate was left open for 10 minutes after the application; once absorption of the applied treatment had taken place the incision was sutured, and the animals kept under observation until they recovered from anesthesia. Once awakened, animals were returned to their tanks in the animal facility.

### All-trans retinoic acid application

Immediately after the right optic nerve was crushed, a strip of Parafilm was placed under it and 10 µl of all-trans retinoic acid (ATRA) solution at 0.1 µM or vehicle (phosphate-buffered saline, PBS 0.1 M) was applied to the region between the stumps. The solution was left in place for 10 min, then the Parafilm was removed and the palate sutured. This dosage and procedure were similar to those used in a previous study [22].

### Resin embedding for light microscopy and transmission electron microscopy

Tissue was embedded in resin as previously described [22]. Euthanasia was carried out one or two weeks after the optic nerve crush and ATRA or PBS application (N=4 per treatment), by using an overdose of anesthetic (1% tricaine +0.04% NaHCO_3_). The optic nerve was dissected from the head and fixed overnight at 4°C in 2% paraformaldehyde + 2% glutaraldehyde, diluted in 0.1 M cacodylate buffer containing 0.05% CaCl_2_. Once fixed the nerve was washed with 0.1M cacodylate buffer, then post-fixed with 1% osmium tetroxide (OsO_4_) in cacodylate buffer for 1 hour under a fume hood. The tissue was dehydrated using 5 minute washes of 25%, 50%, and 70% ethanol, then placed in 3% uranyl acetate diluted in 70% ethanol for 1 hour. Dehydration continued with 90% ethanol and three washes with 100% ethanol (20 minutes each) and a propylene oxide wash for 10 minutes. Finally, the optic nerves were placed in a 50:50 solution of Epon-Araldite and propylene oxide for another hour before leaving them in 100% Epon-Araldite overnight in a dessicator. The next day, the nerves were placed in cubic molds and embedded in 100% resin, then placed in the oven at 60° for 24 hours. The resin blocks were trimmed and, using a Sorvall MT-2 ultramicrotome, cut in semithin sections (1 µm) for light microscopy and ultrathin (90 nm) sections for TEM. Sections were taken from the lesion and regenerating areas. For light microscopy, the semithin sections were stained using methylene blue-azure II and basic fuchsin. For transmission electron microscopy, ultrathin sections were collected on a copper grid and stained using 3% uranyl acetate and 0.075% lead citrate. Imaging and visualization of the ultrathin sections was carried out using a JEOL JEM1011 electron microscope equipped with a Gatan digital camera model 832J46WO.

### Light microscopy cell counts

As in our previous study [22], we cut a series of transverse 1 µm resin sections of optic nerves from distal stump towards proximal stump until the injury site was reached, collecting sections every 50 µm. Analysis was carried out on sections taken approximately 100 µm distal to the injury site, based on previous work showing that regenerating axons have reached this point at one week, and that the number of cells in this region is representative of the cell numbers overall [21]. High magnification light microscope images of the nerve were taken with a CoolSNAP HQ2 camera (Photometrics, Tucson, AZ) and composited using Adobe Photoshop or GIMP. Macrophages were identified based on their large size and dark cytoplasm with granules or vacuoles. It is possible that some of the smallest profiles fulfilling these criteria correspond to phagocytic microglia that are intrinsically in the nerve, therefore we did not include any cells with a diameter smaller than 7 µm. Cell counts were made blind using codes for the image filenames. A color overlay was made outlining the nerve and macrophage-like cell bodies within it; these overlays were subsequently saved as separate grayscale images. ImageJ (FiJi) was used to threshold the grayscale images and to quantify the cell numbers and sizes using the Analyze Particles function.

### Electron microscopic analysis of phagocytic structures

As before, from electron micrographs we quantified sub-cellular structures (“organelles”) indicative of phagocytic activity in macrophage cell profiles [22]. We classified these organelles into three categories: (1) phagocytic vacuoles: electron-lucent, membrane-bound ‘holes’ containing evident cellular debris, the diameter of which measured from 1 to 6 μm; (2) multilamellar bodies, composed of multiple electron-dense lamellae or vesicles, ranging in diameter from 0.5 to 3 μm; and (3) lipid droplets, which ranged from 0.3 to 2 μm in diameter, were generally homogeneously electron-lucent and had no bounding membrane. Color overlays were made in Photoshop or GIMP of 24-40 cells and their phagocytic organelles from 3 animals per experimental group, then these were converted to grayscale images. These were thresholded, and the areas of each organelle type measured using the Analyze Particles function of FiJi. In order to standardize variations caused by partial image cropping, different cell profile sizes, and oblique sections, the total area of each type of structure per cell was expressed as a percentage of the total area of all the organelles. Data were plotted as kernel density (“violin”) plots with superimposed box-whisker plots showing the median and 25–75 percent quartiles (box) and minimum/maximum values (whiskers). Since data were generally not normally distributed, the statistical significance of differences between medians was determined using Kruskal-Wallis tests with *post-hoc* Dunn pairwise comparisons using Bonferroni corrected p-values (*P<0.05, **P<0.01, ***P<0.001).

### Immunohistochemistry

The right optic nerves were dissected while still attached to the eye cup and fixed with 2% paraformaldehyde solution for 30 minutes at room temperature. Then they were washed using 0.1 M PBS, and subsequently placed in 30% sucrose for cryoprotection at 4°C overnight. Next day they were frozen for the subsequent preparation of 12-14 µm thick cryostat sections, which were collected on microscope slides. Prior to immunostaining, the slides were hydrated with 0.1 M PBS for 30 minutes, followed by antigen retrieval using a 10 mM citrate buffer (pH 6) for 10 minutes at 60°C. A pre-incubation with 10% normal goat serum in 0.1M PBS for 1 hour was carried out, following which the slides were incubated overnight at room temperature with the primary antibodies, which were anti-Arginase 1 (1:50; catalog # sc-271430; Santa Cruz Biotechnology), anti-CD68 (ED1, 1:200; catalog # ab125212, Abcam) and anti-Iba1 (1:100; catalog # 01919741; Wako), diluted in 0.1M PBS + 0.3 TritonX-100 + 0.5% BSA. After the washes with PBS 0.1M, the sections were incubated with secondary antibody goat anti-mouse Cy2 (1:200, Jackson ImmunoResearch Laboratories, Inc.) or goat anti-rabbit Cy3 (1:200; Jackson ImmunoResearch Laboratories, Inc.) for 2 hours at room temperature. Sections were then stained using 4’,6-diamidino-2-phenylindole (DAPI) 40µg/mL for 5 minutes at room temperature, washed with 0.1M PBS 3 times 5 minutes each, and finally mounted with Polyaquamount.

Omitting the primary antibodies resulted in the absence of immunostaining. The ED1 antibody recognizes rodent and bovine CD68 or macrosialin, which has some homology with amphibian lysosomal proteins. The Arg1 polyclonal antibody recognizes the C terminus of mammalian arginase 1, the likely antigenic regions of which show almost 70% identity with amphibian arginase. The Iba1 polyclonal antibody recognizes the C terminus of rabbit Iba1 (or “allograft inflammatory factor 1”), a widely used marker for microglia that has been used in a range of species including zebrafish [29]. Because we cannot be completely sure that the antibodies indeed recognize frog homologs of these mammalian proteins, we refer to the staining as “ED1-like immunoreactivity” (ED1-LI), “Arg1-like immunoreactivity” (Arg1-LI) and “Iba1-like immunoreactivity” (Iba1-LI).

### Clodronate liposomes for macrophage depletion

In order to evaluate the importance of macrophages during recovery from optic nerve injury, their numbers were depleted using clodronate liposomes, a technique that is well-established for *in vivo* ablation, and which has been shown to work in zebrafish and frogs [30–32]. We used the Clodrosome® Fluorescent-DiI Macrophage Depletion Kit (Encapsula Nano Sciences, Brentwood, TN), which includes a suspension of 1.5-2 µm liposomes consisting of non-toxic L-alpha-phosphatidylcholine and cholesterol, labeled with 1,1’-Dioctadecyl-3,3,3’,3’-Tetramethylindocarbocyanine (DiI) and containing either disodium Dichloro-phosphono-methyl)phosphonate (Clodronate 18.4 nM) or PBS. 10 µl of liposome suspension were applied to the optic nerve at the time of injury in the same manner as ATRA (see above). When both ATRA and clodronate were applied, the ATRA was removed after 10 minutes and then the clodronate suspension added for a further 10 minutes. Because macrophage repopulation can occur after about a week [31], a second liposome injection was made into the region of the injury site at 1 week after nerve crush.

### Measurements of axonal number

As in a previous study, frozen longitudinal sections of the whole optic nerve were processed with GAP-43 monoclonal antibody (1:500; Chemicon Millipore) in order to stain regenerating axons, allowing us to count the number of axons that projected beyond the injury site [21]. Alternating longitudinal sections through the nerve were analyzed to avoid overlap of data. Axonal regeneration was measured from confocal images obtained with a Nikon laser scanning confocal microscope, using NIS Elements Browser Software. Data was analyzed statistically with ANOVA and *post-hoc* Tukey tests and differences were considered significant if the p-value < 0.05.

## Results

### ATRA application increases the number and size of macrophages 1 week after optic nerve injury

We used light microscopy of transverse 1 μm sections of optic nerve to quantify phagocytic cells. Based on our previous findings (Blanco et al., 2019; Vega-Meléndez et al., 2014) showing that regenerating axons had reached 100 - 200 μm distal to the injury by 1 week, we selected this region for quantifying macrophage-like cells. We identified macrophages based on their large size, dark staining, and granular and/or vacuolar inclusions (Fig 1A and C). In PBS-treated controls, from 70 to 105 macrophage cell profiles were counted in this region from overlays (Fig 1B, N=6 animals). Because the cross-sectional area of the nerve varied somewhat between preparations (0.089 – 0.225 mm^2^), as in the previous publication [22] this cell count was converted to cell density, giving a mean of 654 ± 65 cells/mm^2^ (N = 6). At 1 week, treatment with ATRA resulted in a 65% increase in mean cell density, to 1081 ± 56 cells/mm^2^ (N = 5, p = 0.00011, ANOVA with *post-hoc* Tukey test).

**Fig 1.**
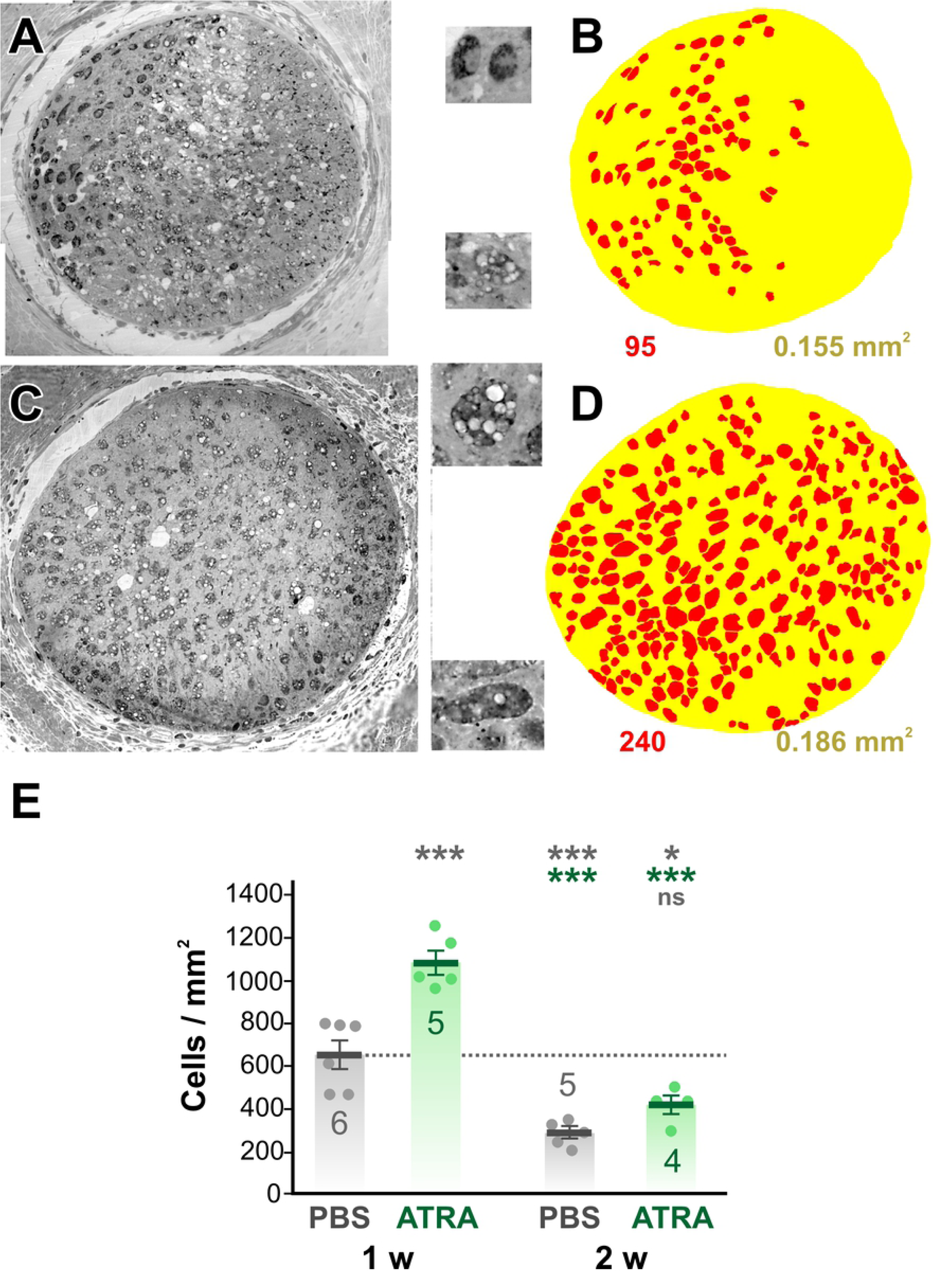
RA application increases the number of macrophages 1 week after optic nerve injury. (A and C) Light micrographs of transverse 1 μm sections of the optic nerve taken from the region 100 - 200 μm distal to the injury area at 1 week after axotomy. A. is a PBS-treated control, C is from an ATRA-treated animal. The insets show enlarged examples of macrophage cell profiles, showing dark staining, granules, and vacuoles. (B and D) Color overlays of the light micrographs, delineating macrophage cell profiles (red) and the nerve itself (yellow). The cell count and nerve area derived from these overlays are shown below. (E) Scatterplots of cell density showing mean ± SEM. Treatments are PBS or ATRA, at 1 or 2 weeks after axotomy. Numbers of animals are shown. Asterisks or “ns” above each column indicate the significance when compared to 1 week PBS (row 1, gray), 1 week ATRA (row 2, red), or 2 weeks PBS (row 3, gray) with ANOVA and *post-hoc* Tukey tests. The macrophage cell density is increased at 1 week after optic nerve injury by ATRA treatment. By the second week the cell density has decreased, with ATRA treatment showing no significant difference from PBS treatment.

At 2 weeks after injury, PBS-treated animals showed a 57% decrease in macrophage cell density compared to 1 week (282 ± 26 cells/mm^2^, N = 5, p = 0.0004). ATRA treatment did not significantly increase the density of macrophages at 2 weeks (417 ± 43 cells/mm^2^, N = 4, p = 0.349). Thus the effect of a single topical application of ATRA is to temporarily increase the numbers of macrophages in the injured optic nerve. When compared to our previous study with application of growth factors, this transient effect of ATRA is comparable to that of FGF-2 application, whereas the effects of a single CNTF application are prolonged until the second week [22].

The macrophage overlays allowed the quantification of various parameters of the cell profiles. We were interested to determine whether the cells became larger as a result of ATRA treatment, and so measured their diameter (Feret diameter, ie. longest diameter of each profile). At 1 week after axotomy, ATRA treatment resulted in a 20% increase in the mean Feret diameter of the macrophages, from 17.7 ± 0.9 μm (N = 6) to 20.4 ± 1.2 μm (N = 5, p = 0.027, homoscedastic t-test). Diameters were segregated in 10 μm bins for each preparation, then these totals were expressed as a percentage of the total number of cells and averaged over the experimental animals (Fig 2). The majority (90%) of the cell profiles fell in the range of 10-40 μm. At 1 week after axotomy, ATRA treatment had significantly decreased the proportion of 10-20 µm diameter cells and increased the number of cells in the 20-30 and 30-40 µm size categories (Fig 2A). No significant changes were seen at 2 weeks in the mean Feret diameter (19.9 ± 2.3 μm, N = 5 vs 19.0 ± 2.1 μm, N = 4) or in the size distributions (Fig 2B).

**Fig 2.**
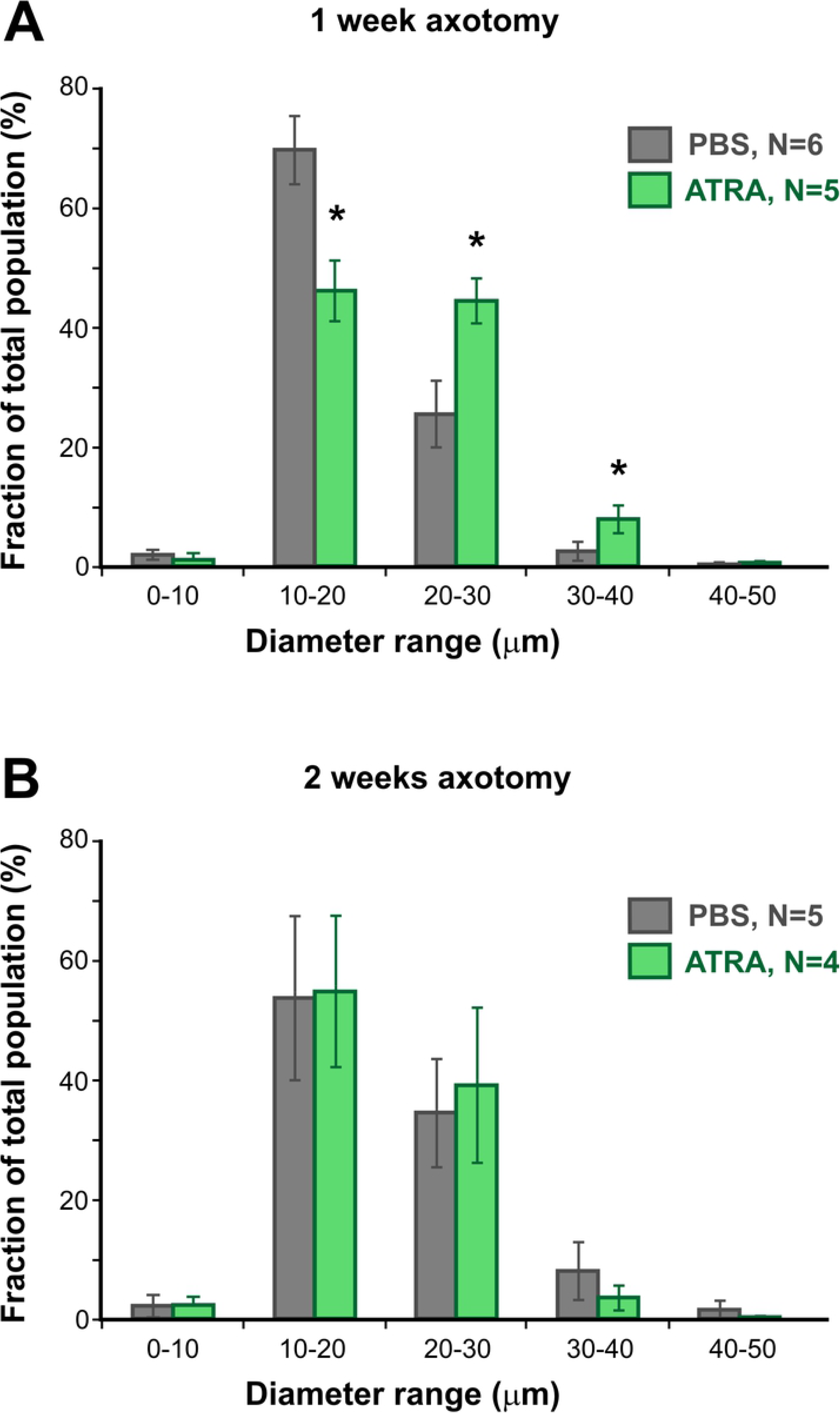
ATRA treatment increases macrophage size at 1 week. (A and B). Frequency histograms of cell profile Feret diameter, averaged over several preparations, showing mean ± SEM for each size category. (A) At 1 week after axotomy ATRA treatment results in a significant decrease in 10-20 µm cells and an increase in 20-40 µm cells (p < 0.05, homoscedastic t-test with sequential Bonferroni correction). (B) No significant changes in cell size are seen at 2 weeks after axotomy

### RA application increases putative microglia at 1 week after optic nerve injury

In our previous study we had noted the presence of microglia in electron micrographs of the regenerating optic nerve, relying on their small size and distinctive morphology to identify them [22]. The question arose as to whether their numbers were increased after axotomy or by treatment with ATRA. In other systems microglia can be distinguished from macrophages by their differing immunoreactivities. The Arg1 antibody recognizes the C terminus of mammalian arginase 1, which is a classic marker for pro-repair M2 macrophages [33]. A BLAST search of the likely antigenic regions of this molecule showed an almost 70% identity with amphibian arginase, and we have shown that it labels macrophage-like cells in frog optic nerve [22]. A commonly used marker for microglia is the polyclonal antibody against C terminus of Iba1 (or “allograft inflammatory factor 1”), which is known to stain zebrafish microglia [29]. We used both antibodies, Arg1 and Iba1, in double-labeling immunostaining experiments to identify putative macrophages and microglia in frozen sections of the injured frog optic nerve at 1 week after axotomy, with or without treatment with ATRA (Fig 3). It appears that small, microglia-like cells that strongly stain with Iba1 are not immunoreactive for the macrophage marker Arg1, and *vice versa* (Fig 3 B, D). In addition, it was found that an increase in Iba1-stained putative microglia was apparent with ATRA treatment. Whereas PBS treatment alone of crushed optic nerve resulted in no significant increase in Iba1-labeled cells per section compared to uncut controls (30.0 ± 10.8, N = 4 vs 24.5 ± 2.3, N = 4), there was a significant 6-fold increase in putative microglial numbers after ATRA treatment (199.5 ± 32.8, N = 4 vs 30.0 ± 10.8, N = 4, p = 0.0005, ANOVA with *post-hoc* Tukey test).

**Fig 3.**
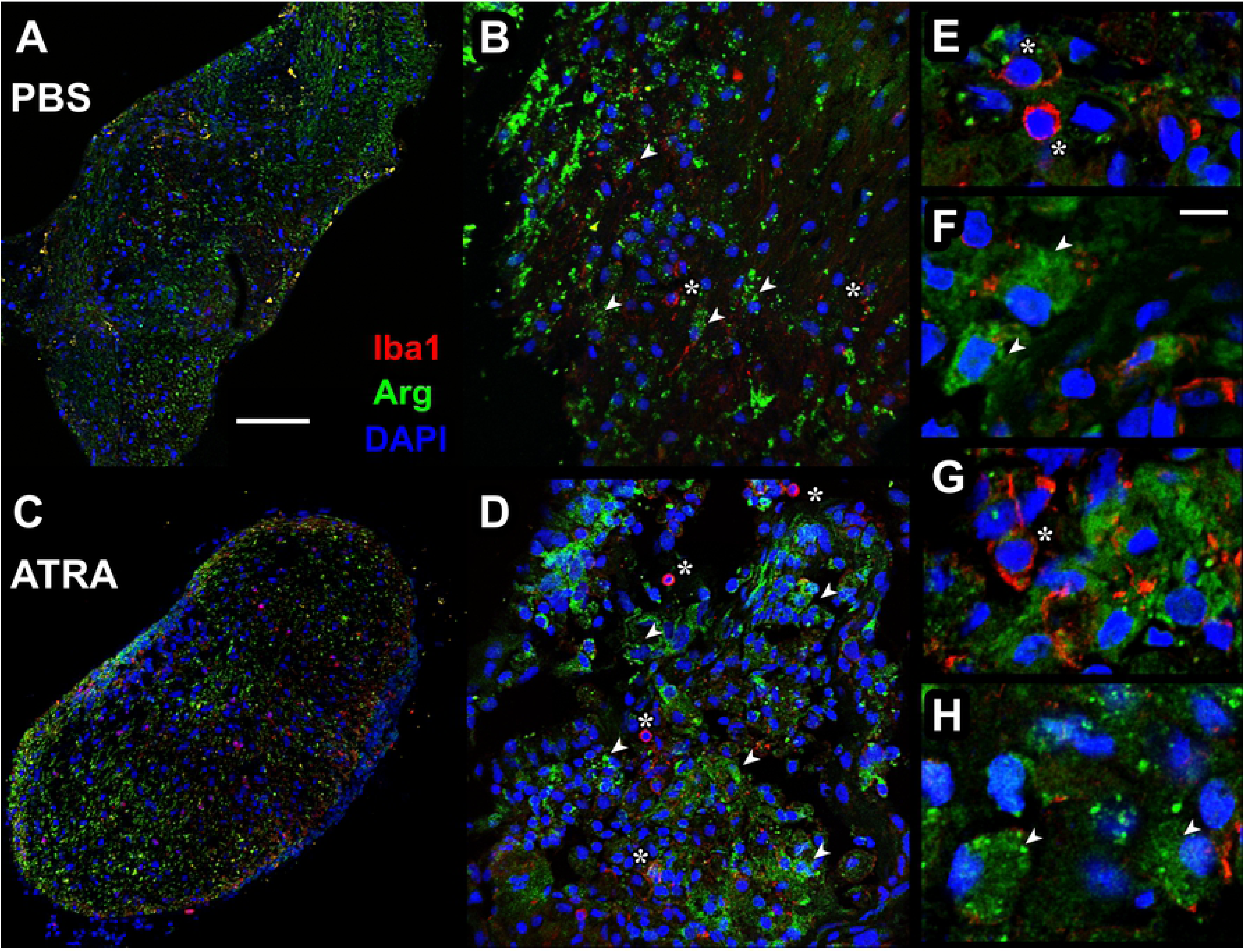
ATRA treatment increases microglia at 1 week. (A - D) Confocal fluorescence micrographs of frozen sections of optic nerve 1 week after axotomy, immunostained with Arg1 (macrophages: green) and Iba1 (microglia: red) antibodies. Nuclei are counterstained with DAPI (blue). (A and C) PBS treatment, (B and D) ATRA treatment, showing increased numbers of Iba1-labeled microglia. (B and D) Higher magnification views with some macrophages (arrowheads) and microglia (asterisks) indicated. Scale bar in A: 100 μm in A and C; 50 μm in B and D.

### Ultrastructural characteristics of macrophages and microglia in the optic nerve after injury

Similar to the previous study [22], at one week after optic nerve crush and PBS treatment actively phagocytosing macrophages were present inside the optic nerve (Fig 4A-D), particularly in the periphery underneath the *glia limitans*, a layer made up of astrocytes, glial cells that characteristically exhibit large bundles of intermediate filaments and desmosomes [34]. Many macrophages contained structures indicative of different stages of phagocytosis (Fig 4 inset). The large phagocytic vacuoles appeared to contain the remnants of degenerating axons and myelin, and presumably indicate the relatively recent occurrence of phagocytosis; multilamellar bodies probably represent a later stage of debris processing; and finally the lipid inclusions are thought to represent the end points of myelin and membrane destruction [35]. At 1 week, there were numerous macrophages with a variety phagocytic organelles (Fig 4D). Two weeks after nerve crush, there appeared to be fewer macrophages in the nerve. Those which were present, both peripherally and centrally, generally contained fewer vacuoles with axonal debris and were predominantly filled with lipid inclusions (Fig 4F and G). A minority of the cells still appeared to be actively phagocytosing at 2 weeks (Fig 4F).

**Fig 4.**
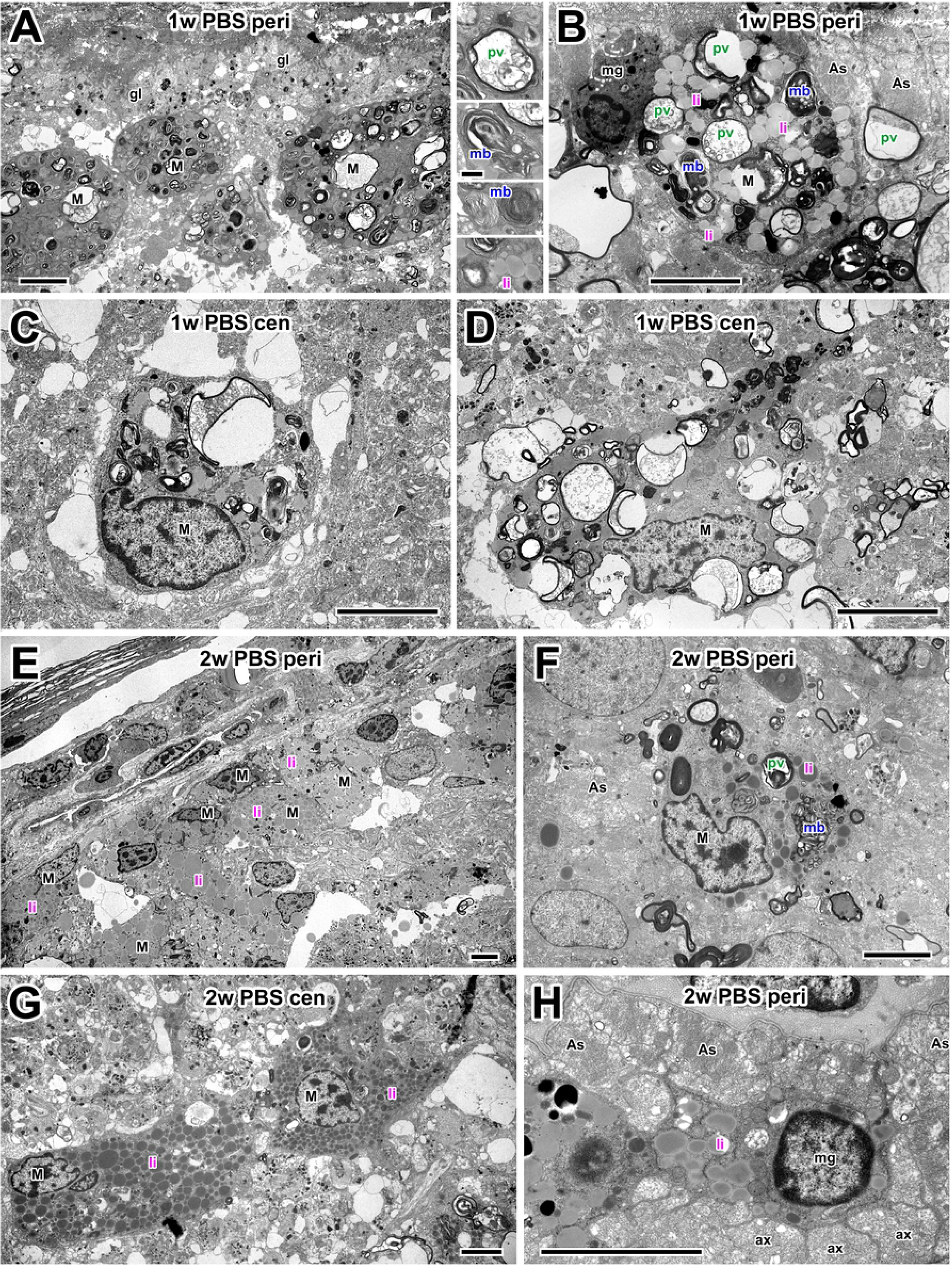
Electron microscopy of macrophages in PBS controls. (A) Medium-low magnification electron micrograph of the optic nerve periphery at 1 w after injury. Macrophages (M) cluster under the *glia limitans* (gl). Inset between A and B shows high-magnification examples of phagocytic organelles: phagocytic vacuoles (pv), multilamellar bodies (mb) and lipid inclusions (li). (B) A peripheral macrophage (M) at 1 w after axotomy containing a mixture of phagocytic vacuoles, multilamellar bodies and lipid inclusions. Next to it is a microglial cell (mg) and astrocytes (As). (C and D) Centrally-located macrophages (M) at 1w, containing a similar range of phagocytic structures and a typical irregular nucleus. (E) At 2 w after axotomy, macrophages (M), predominantly containing lipid inclusions, are clustered at the periphery of the nerve. (F) Peripheral macrophage (M) at 2w, adjacent to astrocytes (As). The macrophage has a variety of phagocytic organelles including phagocytic vacuoles (pv), multilamellar bodies (mb) and lipid inclusions (li). (G) Centrally-located macrophages (M) at 2 weeks after axotomy, containing predominantly lipid inclusions (li). (H) Peripheral microglial cell (mg) containing lipid inclusions (li), adjacent to astrocytes (As) and extending processes between bundles of small regenerating axons (ax). Scale bar: 5 μm in A-H; 1 μm in inset.

In addition to macrophages, smaller cells with evidence of phagocytosis were present, in particular at 1 week, but also at 2 weeks after axotomy (Fig 4B and H). These were identified as microglia by their smaller size, round nucleus, and darker cytoplasm that was typically drawn out into narrow processes that infiltrated between bundles of regenerating axons (Fig. 4H).

At one week after ATRA treatment, more macrophages and microglia were visible in the optic nerve (see quantitative Results sections above). Macrophages showed signs of having been very active in phagocytosis, with many containing predominantly multilamellar bodies (Fig 5A, B). Towards the center of the nerve some showed signs of ongoing phagocytosis, containing phagocytic vacuoles (Fig 5C). Later, at 2 weeks after ATRA treatment, the remaining macrophages were either lipid-filled (Fig 5D, E, F) or still had signs of ongoing phagocytosis (Fig 5E and F). In both 1 week and 2 week preparations many lipid-containing microglia were also present, often near to macrophages and astrocytes (Fig 5E and F).

**Fig 5.**
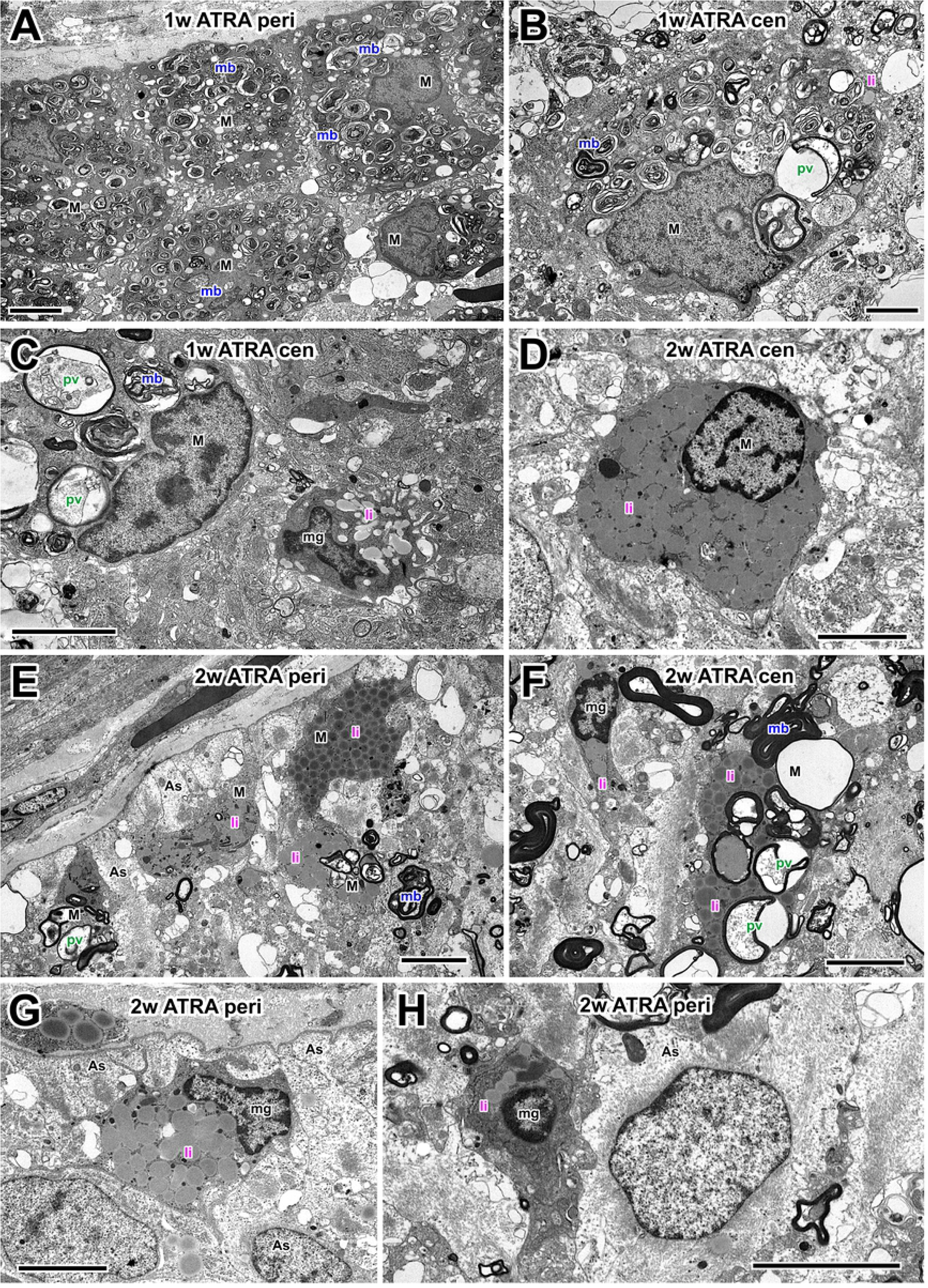
Electron microscopy of macrophages after ATRA treatment. (A) Medium-low magnification electron micrograph of the optic nerve periphery at 1 w after injury, showing several macrophages (M) containing predominantly multilamellar bodies (mb). (B) A macrophage towards the center of the nerve, containing a few phagocytic vacuoles (pv), many multilamellar bodies (mb) and sparse lipid inclusions (li). (C) A macrophage (M) and smaller, putative microglial cell (mg) in the center of the nerve. The macrophage contains phagocytic vacuoles (pv) and multilamellar bodies (mb), whereas the microglia contains predominantly lipid inclusions (li). (D) At 2w after ATRA treatment, a centrally-located macrophage containing only lipid inclusions (li). (E) At 2w after ATRA treatment, macrophage profiles lie below astrocytes (As) at the nerve periphery. The macrophages contain a variety of phagocytic structures. (F) In the central region of the nerve a macrophage (M) with a variety of phagocytic structures, and nearby a microglial cell (mg) with lipid inclusions (li). (G and H) At the periphery of the nerve, microglia (mg) containing lipid inclusions (li) are located between astrocytes (As). Scale bar: 5 μm in A-H.

### Quantitative analysis shows that ATRA prolongs phagocytic activity in macrophages

As in the previous study [22], we carried out a quantitative analysis of the distribution of organelles that are indicative of phagocytosis in macrophages at one or two weeks after axotomy, with or without ATRA treatment. As shown in Fig 6, overlays were constructed from electron micrographs, with color-coding of the three categories of organelle: phagocytic vacuoles, multilamellar bodies, and lipid inclusions or droplets. The large debris-containing phagocytic vacuoles (shown in green) presumably indicate the relatively recent occurrence of phagocytosis; the multilamellar bodies (shown in blue) probably represent a later stage of debris processing; and finally the lipid inclusions (magenta) are thought to represent the end points of myelin and membrane destruction [35]. As before, in order to standardize for variations in profile area due to different cell sizes, oblique sections, and partial image cropping, the total area of each type of organelle per cell was expressed as a percentage of the total area of all the organelles. Again, rather than carry out a multivariate analysis, the phagocytic vacuoles and multilamellar bodies were initially grouped together as “phagocytic organelles”, so as to compare them with the distribution of lipid inclusions (Fig 6C). Finally, we compared the relative contributions by area of the two components of the “phagocytic organelles” group, ie. multilamellar bodies and phagocytic vacuoles (Fig 6D).

**Fig 6.**
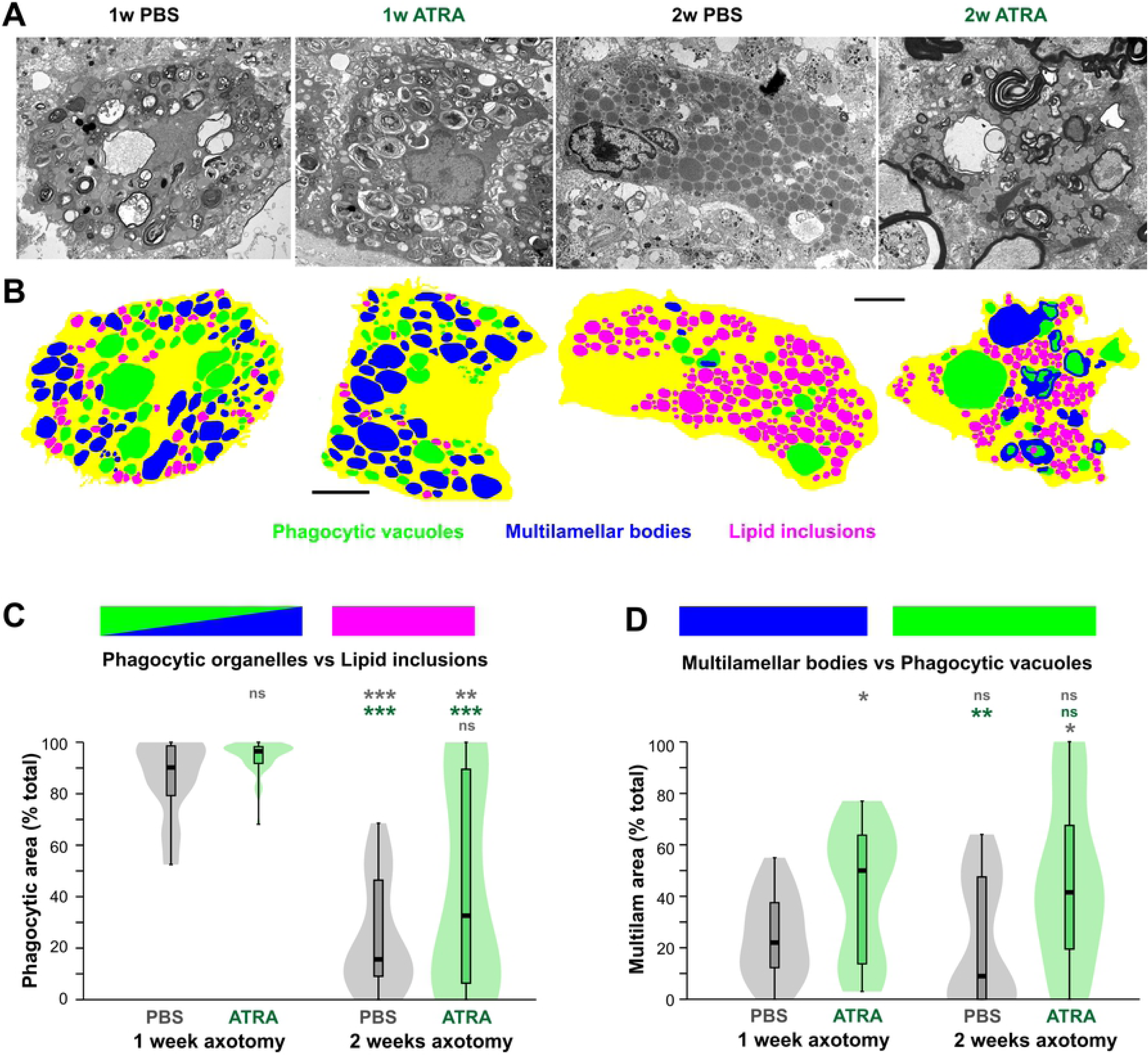
ATRA treatment prolongs phagocytic activity. (A) Representative electron micrographs of macrophages from animals in the four different treatment categories. (B) Color overlays of the cells showing phagocytic vacuoles (green), multilamellar bodies (blue), and lipid inclusions (magenta), with cytoplasm indicated in yellow. Scale bars in A and B: 5 μm. (C, D) Kernel density (“violin”) plots with superimposed box-whisker plots of organelle areas, expressed as a percentage of the total organelle area. Asterisks or “ns” above each column indicate the significance when compared to PBS (rows 1 and 4, gray), or ATRA (row 2 with Kruskal-Wallis test and Dunn’s *post-hoc* comparisons using Bonferroni p-correction (*P < 0.05, **P < 0.01, ***P < 0.001). (C) Comparison of relative areas occupied by “phagocytic organelles” (vacuoles + multilamellar bodies) versus lipid inclusions. At one week, the majority of cells are predominantly occupied by “phagocytic organelles” as opposed to lipid inclusions and ATRA treatment does not change this. At two weeks most cells are occupied predominantly by lipid inclusions in PBS-treated nerves, but there is a significant increase in phagocytic cells with ATRA treatment. N (cells) = 24, 40, 25, 31. (D) Comparison of relative areas occupied by multilamellar bodies versus phagocytic vacuoles. At one week after ATRA treatment, there is a significant increase in the median area occupied by multilamellar bodies. At two weeks, phagocytic vacuoles predominate over multilamellar bodies, but ATRA treatment significantly increases the proportion of multilamellar bodies. N (cells) = 24, 40, 24, 28.

The results of the analysis showed that at 1 week after axotomy, irrespective of ATRA treatment, the great majority of macrophage-like cells contained predominantly “phagocytic organelles” rather than lipid inclusions (Fig 6C). Conversely, at 2 weeks, PBS-treated nerves showed a mixture of cell types, with a large population containing few “phagocytic organelles” and predominantly lipid inclusions, and a smaller population containing more evidence of ongoing phagocytosis. The kernel density (“violin”) plots indicate the possibility of two cell populations, one still undergoing active phagocytosis, and the other with lipid inclusions predominating. Applying an arbitrary cutoff of 50% phagocytic / 50% lipid area, at two weeks 80% of the cells analyzed contained predominantly lipid inclusions, versus 20% with phagocytic organelles. ATRA-treated animals at 2 weeks showed no significant difference in median phagocytic area compared to PBS-treatment (Fig 6C), probably due to the large spread in values. However, in ATRA-treated animals only 58% of the cells had predominantly lipid inclusions, versus 42% with mainly phagocytic organelles, suggesting that ATRA prolongs active phagocytosis.

We then analyzed the relative distribution of the two “phagocytic organelle” types that were previously grouped together; namely, the large debris-containing vacuoles, which presumably indicate the relatively recent occurrence of phagocytosis, and the multilamellar bodies, which probably represent a later stage of debris processing (Fig 6D). At one week after axotomy, ATRA treatment showed a significant difference from PBS controls, with a larger median area occupied by multilamellar bodies compared to phagocytic vacuoles. Using the 50:50 cutoff to define cell populations, in 96% of the cells from PBS-treated animals their phagocytic organelles were predominantly vacuoles rather than multilamellar bodies. On the other hand, in ATRA-treated animals 48% were vacuolar (and 52% multilamellar). At two weeks, 83% of the cells in control animals were predominantly vacuolar, but after ATRA treatment only 64% were. If multilamellar bodies represent a more advanced stage of phagocytic processing, this result again implies that ATRA treatment may accelerate this process.

### Macrophage depletion using clodronate liposomes

In order to be able to test the effects of macrophages on axonal regeneration, we first needed to establish a method for depleting their numbers. Clodronate-loaded liposomes have been shown to be an effective way of doing this, and are effective in zebrafish and frogs [30–32]. At one week after nerve crush and control liposome injection, DiI-labeling could be observed within CD68-immunostained macrophages (Fig 7A). In contrast, one week after injection of clodronate-containing liposomes, both CD68 and DiI staining were largely absent, suggesting that macrophages were absent. Quantitative counts of macrophage-like cells were made from overlays of 1 µm resin sections, as described above (Fig 7C, D). Mean cell density was not significantly affected by PBS-containing liposomes (658 ± 111 cells/mm^2^ versus 654 ± 65 cells/mm^2^ in PBS controls, ANOVA with *post-hoc* Tukey test). However, there was an 84% decrease to 105 ± 40 cells/mm^2^ in clodronate-liposome-treated animals (N = 4, p = 0.0017).

**Fig 7.**
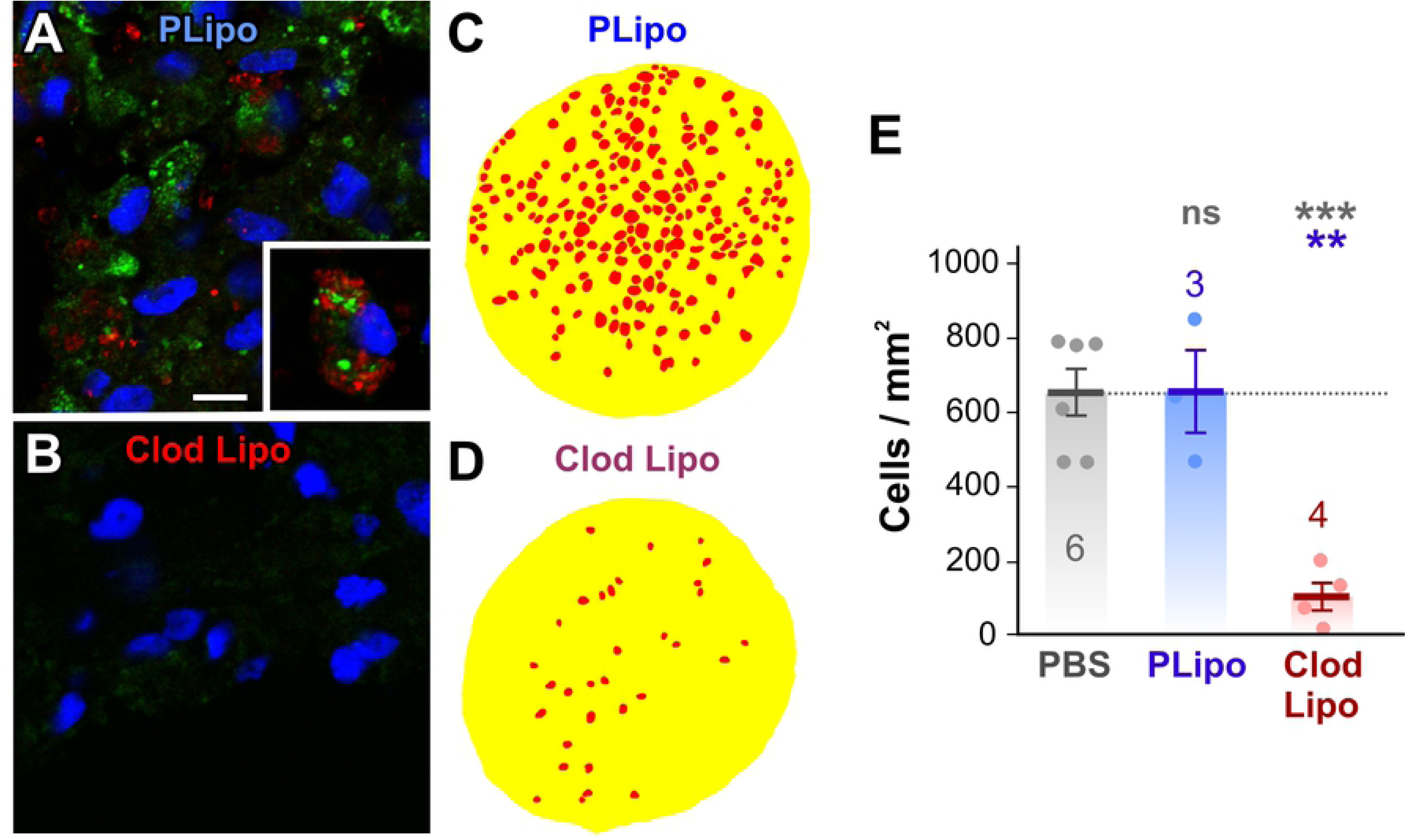
Clodronate liposomes deplete macrophage numbers. (A, B) Immunohistochemical staining of macrophages with anti-CD68 antibody in the optic nerve at 1 week after axotomy and treatment with (A) DiI-labeled liposomes and PBS (PLipo) or (B) with DiI-labeled clodronate liposomes (Clod Lipo). Nuclei are counter-stained with DAPI (blue). (A) Macrophages stained with CD68 (green) are also labeled to varying degrees with DiI-labeled liposomes (red). The inset shows a macrophage with particularly strong liposome uptake. Scale bar: 10 μm. (B) Clodronate-liposome-treated nerve shows cells with only background staining with CD68, and no detectable DiI-liposome labeling. (C, D) Color overlays of light micrographs of transverse 1 μm sections of the optic nerve taken from the region 100 μm distal to the injury area at 1 week after axotomy and liposome treatment, delineating macrophage-like cell profiles (red) and the nerve itself (yellow). (E) Scatterplots of cell density showing mean ± SEM. Treatments are PBS alone, PBS liposomes (PLipo), or clodronate liposomes (Clod Lipo), at 1 week after axotomy. Numbers of animals are shown. Asterisks or “ns” above each column indicate the significance when compared to 1 week PBS control (row 1, gray), or 1 week clodronate liposomes (row 2, blue) with ANOVA and *post-hoc* Tukey tests. The macrophage cell density is unaffected by DiI-labeled liposome treatment alone. However, clodronate liposome treatment resulted in an 80% decrease in macrophage density.

### Numbers of regenerating axons are increased by ATRA and decreased by clodronate

Having established above that ATRA increases macrophage numbers and phagocytic activity, and that clodronate liposomes deplete macrophages by about 80%, we then assayed the effect of these treatments on the numbers of regenerating axons. To this end we used a similar method as reported previously, by immunostaining regrowing axons in longitudinal nerve sections with anti-GAP-43 antibody and counting their numbers per section at a distance of about 200 μm distal to the lesion site, in the regenerating zone [21].

In crushed PBS-treated nerves there were on average 84 ± 3 axons per section (N = 5 animals) (Fig 8F). ATRA treatment increased this number by 53%, to 129 ± 6 axons per section (N = 8, p = 1.7 x 10^-5^, ANOVA with *post-hoc* Tukey test) (Fig 8F). This suggests that (1) retinoic acid could act directly on the axons at the injury site and thereby increase the rate of their regrowth, and/or (2) the increased numbers of phagocytosing macrophages induced by ATRA (see results above) promote the regeneration of the axons.

**Fig 8.**
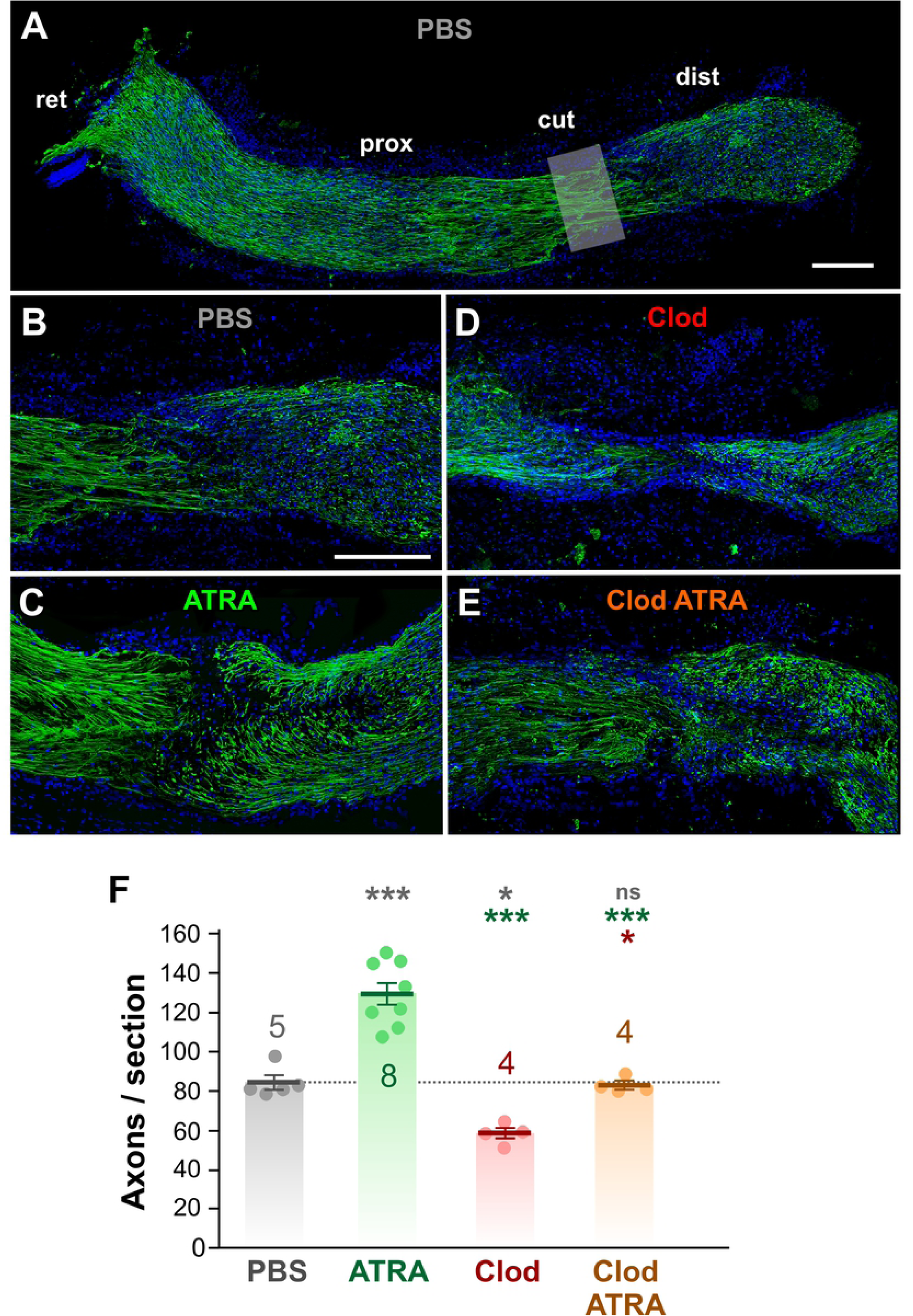
Regenerating axons are increased by ATRA and decreased by clodronate. (A - E) Confocal images of immunohistochemical staining of growing axons with anti-GAP43 antibody (green), in longitudinal sections of the optic nerve at 2 weeks after axotomy. Nuclei are counter-stained with DAPI (blue). (A) Low magnification maximum intensity projection of the complete nerve from a PBS-treated animal, composited from several 12 µm sections, with the retina (ret) on the left and the distal stump (dist) on the right. The cut site (cut) and proximal stump (prox) are also indicated. (B-E) Enlarged views of the cut zone and regenerating regions of nerves from animals with different treatments. (B) PBS treatment, composite from 2 sections (C) ATRA-treatment, single section (D) Clodronate liposome treatment, composite from 2 sections. (E) ATRA and clodronate liposome treatment, composite from 2 sections. Scale bar: 100 μm in A and B. (F) Bar and scatter plots of the number of axons per transverse section, sampled at approximately 200 μm distal to the cut site, showing mean ± SEM. Treatments are PBS alone, ATRA, clodronate liposomes (Clod), or clodronate liposomes along with ATRA (Clod ATRA). Numbers of animals are shown. Asterisks or “ns” above each column indicate the significance when compared to PBS control (row 1, gray), ATRA (row 2, green), or clodronate (row 3, red) with ANOVA and *post-hoc* Tukey tests. Compared to PBS-treated, the number of regenerating axons is doubled by ATRA treatment, decreased by 30% by clodronate, and, in the case of ATRA combined with clodronate, is not significantly different.

In order to test the second hypothesis, clodronate liposome treatment was used to deplete the numbers of macrophages. Clodronate liposome treatment alone decreased the numbers of regenerating axons by about 30% compared to PBS (58 ± 3 axons per section, N = 4, p = 0.0187). In combination with ATRA application, clodronate liposomes completely prevented the ATRA-induced increase in numbers of regenerating axons compared to PBS (Fig. 4F) (83 ± 2 axons per section, N = 4, p = 0.9978). These results support the idea that much of ATRA’s effect on stimulating axonal regeneration is due to the recruitment of macrophages, rather than a direct effect on the axons themselves. The fact that clodronate alone results in 80% macrophage depletion but reduces regeneration by only 30% suggests that the normally-occurring entry of macrophages into the injured nerve is indeed important for axonal regeneration, but also perhaps that other instrinsic factors are in play, which are unaffected by clodronate.

## Discussion

The first result reported in this study shows that application of all-trans retinoic acid (ATRA) at the time of nerve crush results in a 65% increase in the number of macrophage-like cells that enter the regenerating nerve stump at 1 week after injury. When compared to our previous study with application of growth factors, this transient effect of ATRA is most comparable to that of FGF-2 application; in contrast, the effects of a single CNTF application are prolonged until the second week [22]. The magnitude of the ATRA effect is not as large as that of the growth factors, however, unlike the growth factors, ATRA induces a small but significant increase in the cell size.

It has been known for many years that macrophages congregate at the site of nerve injury. Earlier work from this laboratory and others noted the accumulation of macrophages in the axotomized optic nerve of the frog *Rana pipiens* [34, 36], which we later quantified and showed to be preponderantly of the M2 neuroprotective type [22]. Macrophages in other animals show similar large yet transient influxes after nerve injury, for example rat sciatic nerve [37], axolotl spinal cord and peripheral axons [38], and the optic nerves of rat, goldfish and *Xenopus* tadpoles [39–41].

How does ATRA increase the entry of macrophages? The migration of bone-marrow-derived macrophages to the injury site is induced by chemokines, a type of chemoattractant cytokine, secreted by the injured tissue. Of particular importance is monocyte chemoattractant protein-1, now known as CCL2 [42, 43]. Our present finding that ATRA promotes the entry of macrophages suggests that either (1) RA itself can act as a chemoattractant for bone marrow-derived macrophages, or (2) that it promotes the secretion of chemokines such as CCL2 from cells in the injured region such as microglia and astrocytes [44], and perhaps oligodendrocytes.

We have not found any evidence that RA itself acts as a direct chemoattractant for macrophages. Regarding the second hypothesis, that RA stimulates chemokine secretion, there is evidence that it can increase CCL2 expression in, and promote migration of, human monocytes [45]. On the other hand, RA has an anti-inflammatory effect on mouse astrocytes *in vitro*, suppressing the release of CCL2 (and other chemokines) [46]. Retinoic acid-inducible gene I (RIG-I) plays a role in mediating the up-regulation of CCL2 expression by mouse microglia in response to viral infection [47] but it is not known whether RA would have the same effect. Otherwise, this hypothesis remains to be tested.

In the present study we also show that ATRA also results in a large increase in numbers of Iba1-positive putative microglia, a phagocytic cell type that is intrinsic to the CNS. There are few studies that have addressed this question. In one, RA decreases mouse microglial activation *in vitro* [48], this being one suggested mechanism for its neuroprotective effect [49]. On the other hand, RA-loaded nanoparticles do increase the preponderance of the neuroprotective M2 phenotype of microglia in murine hippocampal slices [50].

We then present electron microscopic evidence that ATRA prolongs phagocytic activity in macrophages, similar to the effects of the growth factors CNTF and FGF-2 [22]. This is consistent with recent evidence that RA promotes macrophage phagocytosis of myelin debris. Retinoic acid, acting via RARβ receptors, induces the non-inflammatory M2 phenotype of mouse macrophages *in vitro*, and increases their ingestion of myelin, via the upregulation of genes associated with phagocytosis [27]. Retinoid X receptor activation in aged macrophages reverses their deficiencies in myelin debris phagocytosis and remyelination *in vivo* and *in vitro* [51]. It must be pointed out, however, that the percentage of myelinated axons in the frog optic nerve is low, approximately 3% [34], so it is possible that clearance of debris from degenerating unmyelinated axons is of equal importance in facilitating axonal regrowth.

We then used the well-established method of treatment with clodronate-loaded liposomes to reduce macrophage numbers, first demonstrating that this is 80% effective in the frog optic nerve. Finally, having established above that ATRA increases macrophage numbers and phagocytic activity, and that clodronate liposomes deplete the macrophages, we then tested the effect of these treatments on the numbers of regenerating axons. We show that, in a manner similar to CNTF and FGF-2 [21], ATRA increases the numbers of regenerating axons in the distal stump of the optic nerve by 53%.

In principle this could have been due to a direct effect of ATRA on axonal growth since RA, acting via the RARβ2 receptor, promotes neurite outgrowth from cultured mouse spinal cord [52], and RARβ2 overexpression increased regeneration in rat spinal cord [53]. We have shown that RARβ immunoreactivity is present in regenerating frog RGC axons [19]. RA promotes and direct neurite outgrowth of invertebrate neurons via effects on Ca^2+^ signaling [54].

While we cannot rule out the possibility that ATRA directly increases RGC axon growth, our results show that clodronate liposome treatment was sufficient to completely prevent the ATRA-induced increase in numbers of regenerating axons. Clodronate also prevented the increase in numbers of macrophages, suggesting a direct causal relationship between the numbers of phagocytosing macrophages and axonal regrowth. In support of this, we also found that depletion of macrophages alone, independently of ATRA treatment, reduced the numbers of regenerating axons by 30%. In contrast, in mouse optic nerve, depletion of macrophages alone had no effect on lens-injury-induced axonal regeneration [55]. This may perhaps be in part due to the high M1/M2 macrophage ratio observed in mouse CNS, which may not be conducive to regeneration [56], whereas in amphibia we have shown that the M2 pro-regenerative type may be predominant [22]. In some cases there is a direct link between myelin debris removal and axonal regeneration, for example, in mouse peripheral nerve injury, doubling the rate of myelin phagocytosis (via SIRPα deletion) increases the removal of myelin debris and doubles the numbers of regenerating axons [57].

In conclusion, our results show that ATRA treatment of the injured frog optic nerve increases the numbers of M2-type macrophages and putative microglia, and increases the amount of phagocytosis by those macrophages. The molecular mechanisms behind these effects remain to be determined. ATRA treatment also increases the numbers of regenerating axons in the distal nerve stump. Clodronate depletion experiments suggest that the success of axon regeneration is partially dependent on the presence of debris-phagocytosing macrophages, and that the increases in axonal regeneration caused by ATRA are primarily due to the increased numbers of macrophages that it induces, rather than being a direct effect. Further studies will examine whether macrophage depletion also affects RGC survival.

## Acknowledgments

The authors wish to gratefully acknowledge the contribution of the late Clarissa Del Cueto, who will be greatly missed. Without her dedication and technical expertise much of this work would not have been possible.

## Supporting Information Captions

**S1 Table.** Experimental data from light microscopy cell counts, including the results of ANOVA and Tukey tests.

**S2 Table.** Experimental data from counts of GAP-43-stained regenerating axons, including the results of ANOVA and Tukey tests.

## Notes

### Competing Interest Statement

The authors have declared no competing interest.

